# SAGA (Simplified Association Genome-wide Analyses): a user-friendly Pipeline to Democratize Genome-Wide Association Studies

**DOI:** 10.1101/2025.08.25.672146

**Authors:** Basilio Cieza, Neetesh Pandey, Vivek Ruhela, Sarwan Ali, Giuseppe Tosto

## Abstract

Genome-wide association studies (GWAS) have enabled clinicians and researchers to identify genetic variants linked to complex traits and diseases(1–3). However, GWAS still face several challenges, particularly regarding accessibility and reproducibility (4–6). Conducting these analyses often requires substantial bioinformatics expertise for data preprocessing, software installation, and scripting(7–10). We then developed SAGA (“*Simplified Association Genome-wide Analyses”*), a BASH-based, open-source, fully automated pipeline that integrates three widely adopted tools—PLINK(11), GMMAT(12), and SAIGE(13)—for accessible, robust, and reproducible GWAS. After installation, users simply need to provide genotype and phenotype files in standard formats. The pipeline automates preprocessing, association testing, and visualization, outputting summary statistics, Manhattan plots, and quantile-quantile plots. SAGA enables robust GWAS for users without scripting experience, expanding access to complex genetic analyses.

## 1. Introduction

Genome-wide association studies (GWAS) have transformed discovery of gene-variant associations for complex traits and diseases in human populations (14–16). Yet, the broader deployment of GWAS is limited by technical barriers: multi-step software installations, data formatting, and intricate analytic workflows (17–19). SAGA (“*Simplified Association Genome-wide Analyses”*) addresses this gap for non-bioinformaticians with a modular, bash-based pipeline that integrates three popular platforms: **PLINK** for standard linear/logistic regression (11), **GMMAT** for generalized mixed model association testing, suited for related or family-based samples (12), and **SAIGE** for large, imbalanced case-control studies and rare variant analysis (13). SAGA offers an automated, reproducible solution for GWAS analysis on mainstream genomic data, providing ready-to-use outputs for downstream exploration and publication.

## 2. Methods

### Tool Selection and Rationale

We implemented PLINK, GMMAT, and SAIGE for their complementary strengths (11–13) (**Table 1**).

**Table 1.** Software included in SAGA and their features.

| **Tool** | **Key Strengths** | **Ideal For** | **Model Type** | **Handles Relatedness** | **Handles Case-Control Imbalance** |
| --- | --- | --- | --- | --- | --- |
| **PLINK** | Fast, user-friendly | Unrelated individuals | Linear/logistic regression | NO | NO |
| **GMMAT** | Mixed models, kinship-aware | Family/related samples | Linear/logistic mixed model | YES | NO |
| **SAIGE** | Scalability, rare variants | Biobank, unbalanced traits | Saddlepoint Approximation (SPA) | YES | YES |

### Pipeline Workflow

The user simply has to shape input genetic files in traditional PLINK’s binary format (.*bed/*.*bim/*.*fam*) along with a structured phenotype file containing family and individual IDs (“*FID*” and “*IID*”, respectively), outcome measures (binary or quantitative), and optional covariates. All input files undergo automated validation to ensure completeness and correct formatting (**Supplementary Figure 1** for more details). In the preprocessing stage, SAGA performs quality control (QC) to filter variants and samples based on missingness, Hardy–Weinberg equilibrium, and minor allele frequency (MAF) thresholds as best practice guides suggest (20–22) (**Supplementary Table 1** and **2**). Population structure is assessed via principal component analysis (PCA), generating the top ten principal components for covariate adjustment (23,24) using PLINK. To account for relatedness in GMMAT, SAGA generates a dense genetic relationship matrix (GRM) using GEMMA (25); on the contrary, SAIGE builds internally a sparse GRM which will be used in subsequential steps. Each preprocessing step is standardized and error-checked to minimize user intervention. Association analysis is conducted using one of three backends, selected automatically according to user-defined parameters or data characteristics: PLINK for fast linear or logistic regression on no related populations, GMMAT for generalized linear mixed models in related populations, and SAIGE (which uses a saddlepoint approximation “SPA”) for mixed model designed to handle large-scale datasets (e.g. biobanks) featuring highly unbalanced case-control ratios and analyses of rare variants. Finally, post-analysis procedures standardize summary statistics across backends and produce publication-ready Manhattan and quantile-quantile (QQ) plots in both high-resolution .*png* and .*pdf* formats using automated R scripts (26). This fully integrated workflow ensures consistent, reproducible results from raw genotype data to final visualization.

### Technical Implementation

SAGA is implemented entirely in BASH and requires only a UNIX/Linux environment for execution. The pipeline depends on PLINK for genotype processing, R 4.2.2 for running GMMAT and generating plots, and Singularity container of SAIGE (version 1.3.0) for scalable mixed-model association testing. Installation is straightforward via a single command:

*git clone* https://github.com/bciezah1/SAGA.git

Upon completion of the analysis, SAGA generates all results in an organized output/ directory. This includes standardized summary statistics (“*summary_stats*.*txt*”) and high-quality plots (Manhattan, QQ, and SNP density plots).

### Automated Preprocessing Details

Once installed, the package can be launched via a single command.

For example, employing PLINK with a binary phenotype and three covariates:

.**/run_pipeline_plink.sh \**

*{user’s path}***/toy_data/geno \**

*{user’s path}***/toy_data/pheno_binary.txt \**

**COV1, COV2, COV3, PC1, PC2, PC3 \**

**PHENO \**

**binary \**

**OutputFolderLabel**

In this example, the second argument specifies the location of the genotype data in PLINK format, followed by the phenotype file location, a comma-separated list of covariates (“*COV1, COV2, COV3*”) and principal components to include (“*PC1, PC2, PC3*”), the target phenotype name (“*PHENO*”), the phenotype type (“*quantitative”* or “*binary*”), and the label of the folder where the results will be located (“*OutputFolderLabel*”). Upon completion, SAGA generates standardized summary statistics (“*summary_stats*.*txt”*) and high-quality Manhattan and QQ plots (“*manhattan*.*png” and “qqplot*.*png”* (**Supplementary Table 2** for additional details).

## 3. Benchmarking

### Simulated Data

We benchmarked the performance and usability of SAGA, using in-house simulated datasets that varied in complexity in a controlled manner. Specifically, we simulated datasets with three different sample sizes (N=1,000, 5,000, and 10,000 individuals), with 100 000 single nucleotide polymorphisms (SNPs; **Figure 1, Supplementary Table 3**). Phenotypes and covariates were simulated to mimic binary traits with typical covariate structures encountered in common GWAS studies. Each dataset was analyzed separately using the three GWAS analysis backends integrated within SAGA.

**Figure 1.**
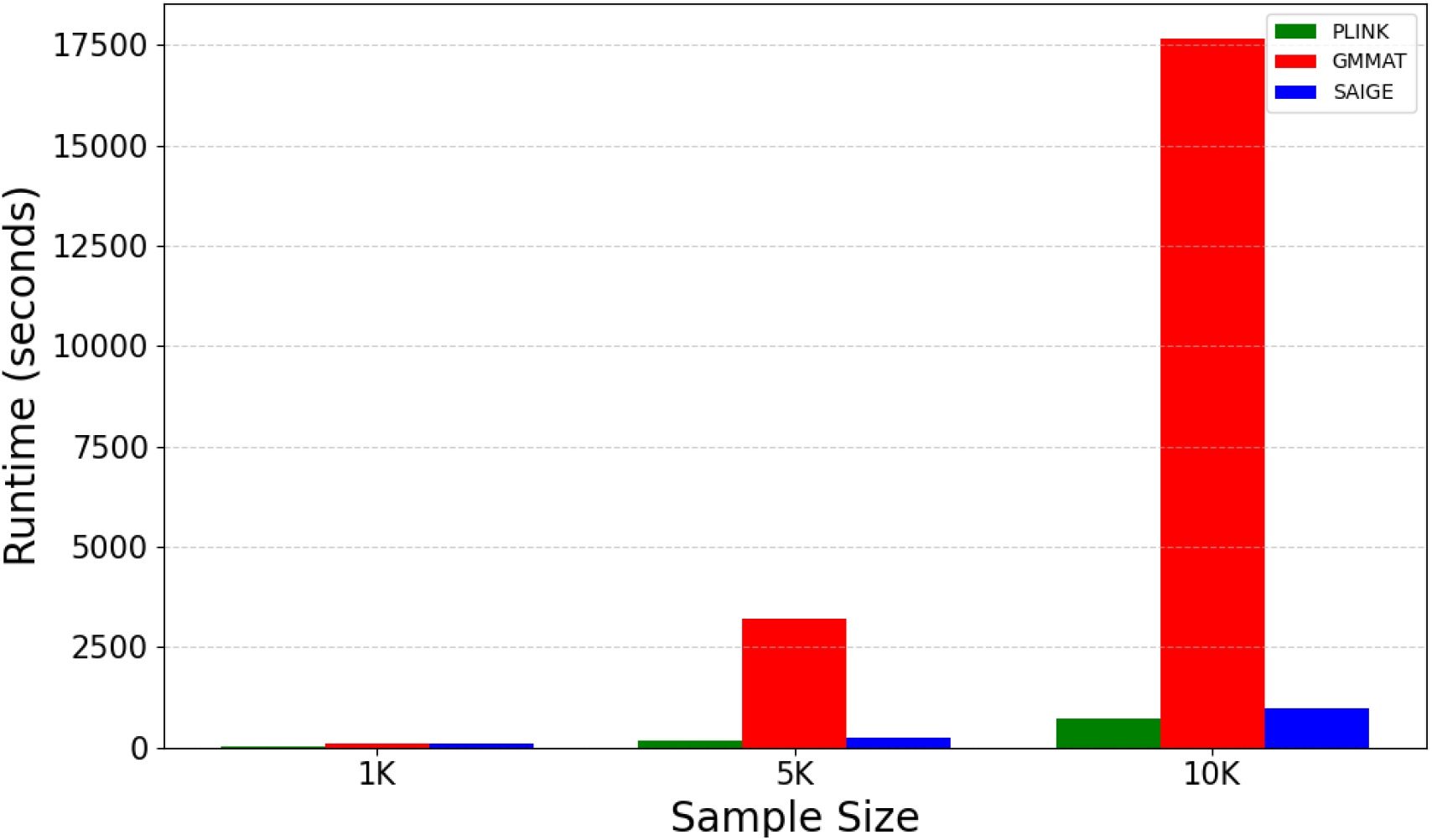
Benchmark for three different samples sizes and 100,000 SNPs using PLINK, GMMAT, and SAIGE (green, red and blue color, respectively).

### Real Data

We further evaluated the performance and applicability of SAGA using real data from the Alzheimer’s Disease Sequencing Project (ADSP) (27). Specifically, we analyzed seven independent cohorts — ADC, PeADI, NOMAS, PRADI, PR1066, TARCC, and WHICAP — each representing diverse populations and study designs.

For each cohort, genotype and phenotype data were processed independently through the full SAGA workflow, including sample and variant QC, SNP pruning, covariate adjustment, and genome-wide association testing. The primary phenotype, Alzheimer’s disease (AD), was modeled as a binary trait, with age, sex, and the top five ancestry-derived PCs included as covariates.

Each cohort was analyzed individually using SAGA (PLINK, GMMAT, and SAIGE. In addition, all cohorts were combined into a single multi-sample to perform a joint genome-wide association analysis, enabling us to test SAGA under large sample sizes (in addition to increased statistical power and cross-cohort consistency assessment). Results for the ADSP multi-sample using SAIGE are presented in **Figure 2** and **3** (**Supplementary Figure 3** and **4** for PLINK and GMMAT results, respectively), while results from individual cohort are provided in **Supplementary Figures 4-10**.

**Figure 2.**
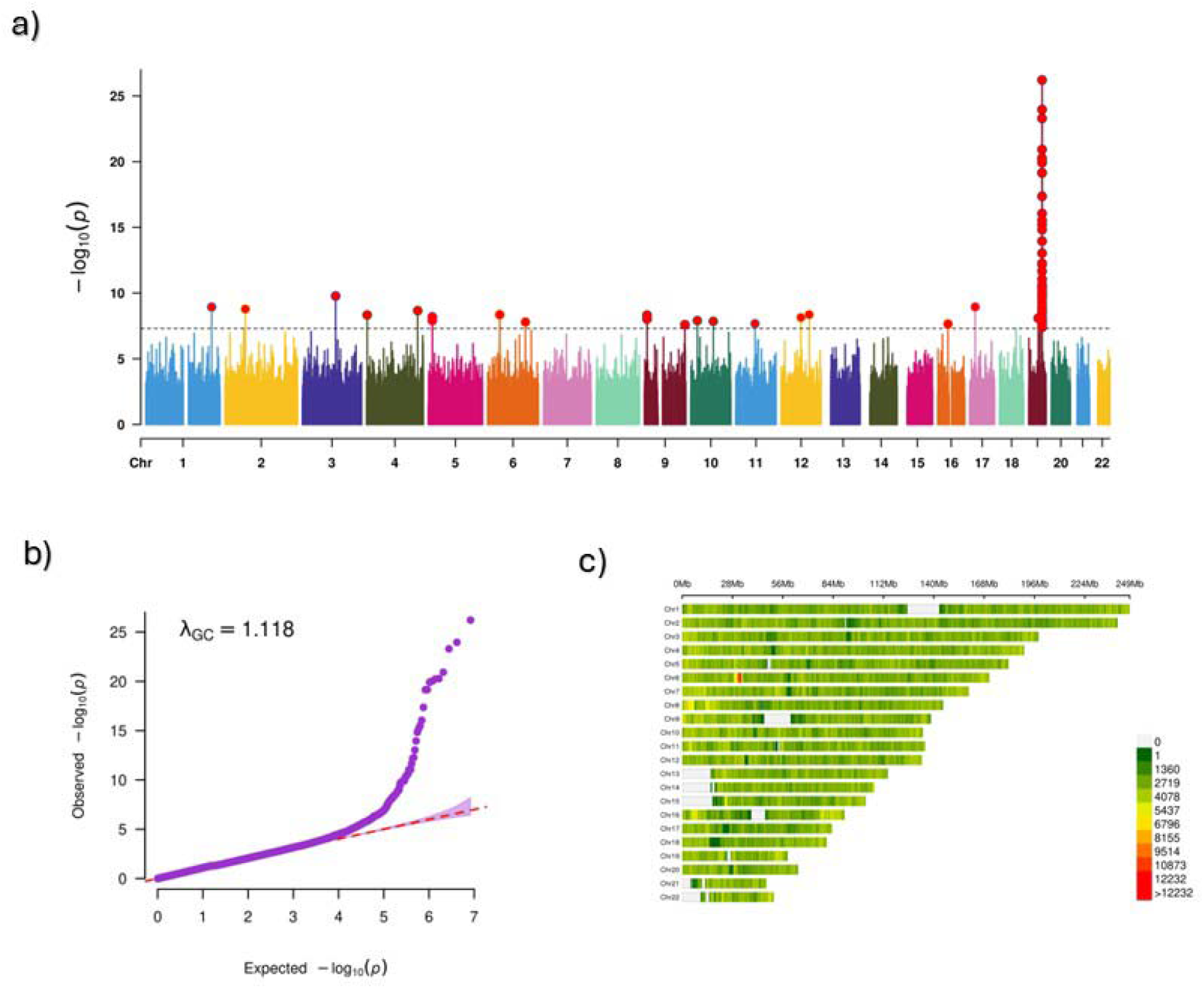
Manhattan plot, QQ, and SNP density plot for common variant analysis using SAIGE.

**Figure 3.**
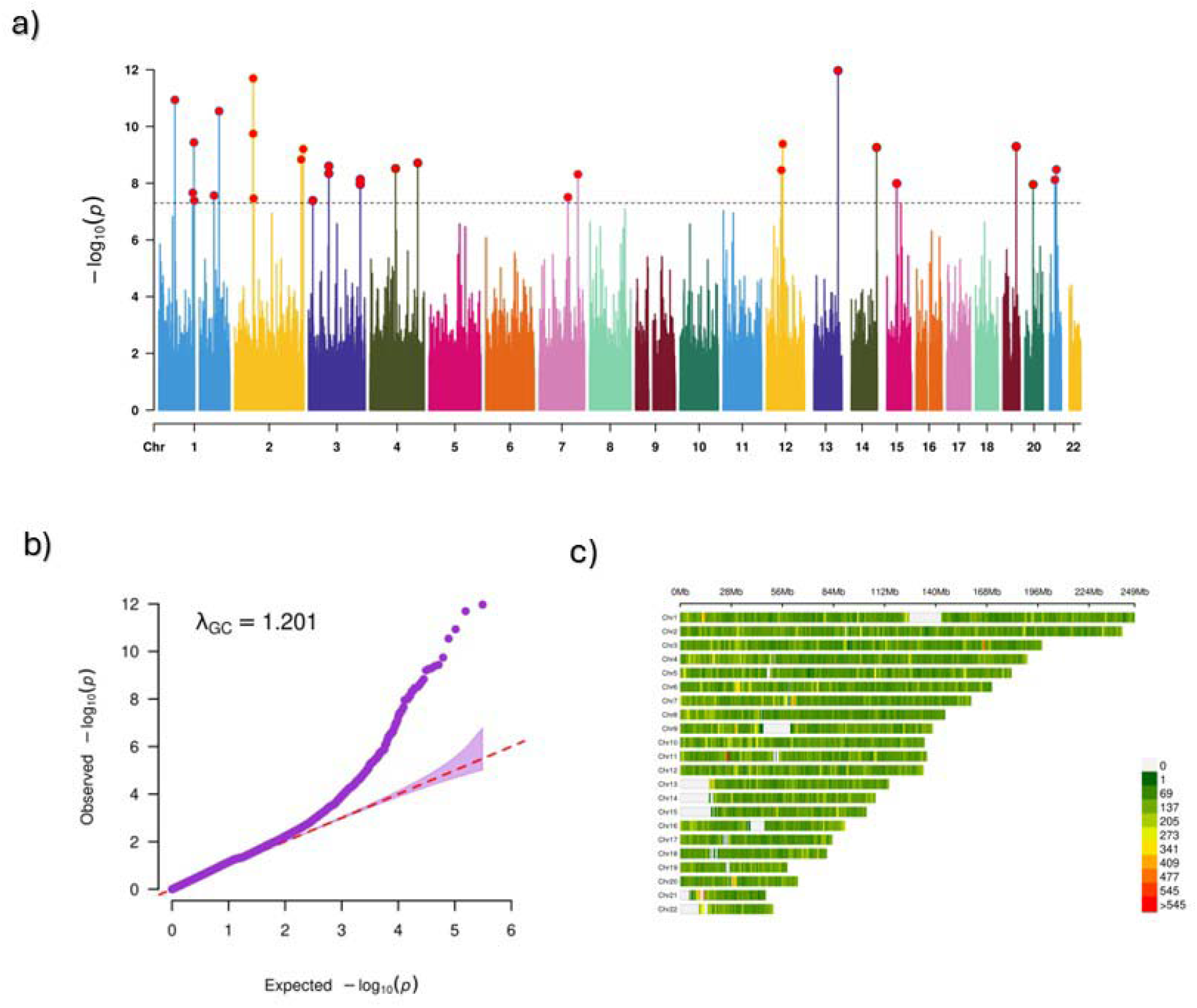
Manhattan plot, QQ, and SNP density plot for rare variant analysis using SAIGE.

## 4. Discussion and conclusions

SAGA provides GWAS workflow for users without scripting or programming expertise by offering an intuitive and automated framework. One of its major advantages is ease of access, as users do not need to edit code and only require minimal familiarity with command-line operations (customized coding is of course optional to the user’s end). The pipeline covers all critical steps of genome-wide association studies, from quality control through association analysis to the generation of publication-ready plots, thereby significantly reducing the technical burden on researchers. By employing consistent, standardized processing methods compliant with leading best practices (20–22), SAGA ensures reproducible results across multiple runs. Importantly, it includes rigorous quality control measures that account for population structure and relatedness among samples, facilitating robust association testing.

Our benchmarking results highlight the strengths of SAGA in terms of runtime efficiency, error control, reproducibility, and usability. Moreover, SAGA substantially simplifies GWAS implementation, enabling researchers - even those with limited computational experience - to perform large-scale analyses quickly and reliably as we showed in our analysis of ADSP data. Importantly, the framework demonstrated the ability to scale with increasing dataset sizes and variant complexity while preserving automated and reproducible workflows across multiple statistical models and software implementations. These findings underscore the utility of SAGA for a wide range of GWAS applications and emphasize its potential to broaden access to advanced genetic association analyses for researchers without extensive computational expertise.

SAGA has some limitations. It assumes that users provide valid and properly formatted phenotype files, as it does not incorporate functionality to check or compose phenotypic data. Additionally, SAGA currently lacks native support for meta-analysis or the merging of multiple cohorts, which are important considerations for large-scale genetic studies. While the pipeline can handle moderately large datasets, analyzing very large datasets—such as those exceeding 50,000 individuals—may still require high-performance computing clusters or cloud resources due to memory and processing demands. SAGA has been tested and optimized for Linux-based systems and has not yet been validated on Windows or macOS environments. Moreover, the SAIGE component is executed through a Singularity image, which requires a compatible cluster or server environment and is not intended for use on local workstations or laptops.

We have plans to include the addition of modules for gene-based rare variant analyses, which would extend the pipeline’s scope to capture a broader spectrum of genetic variation. The developers also intend to implement advanced post-GWAS annotation functionalities, including compatibility with tools for pathway analyses, which can facilitate the biological interpretation of significant loci. Another anticipated feature is support for polygenic risk scores, enabling the translation of GWAS findings into quantitative measures of genetic risk for complex diseases. These enhancements aim to further empower researchers with limited computational backgrounds to perform comprehensive and insightful genetic association studies.

## DATA AND SOFTWARE AVAILABILITY

All code, documentation, and test data are available at: https://github.com/bciezah1/SAGA.git

## AUTHOR CONTRIBUTIONS

**BC, GT:** Conceptualization, manuscript writing

**BC:** software development

**BC, NP, VR, SA, GT:** documentation, testing

## ETHICAL STATEMENT

No human or animal subjects were involved in this work. All testing datasets were simulated and fully anonymized.

## ACKNOWLEDGEMENTS

We thank the GiusTo Lab at Columbia University for cohort data access and deployment support. This work is supported by NIA/NIH award U01AG081817, RF1AG082009 and U19AG074865.

## Notes

https://github.com/bciezah1/SAGA.git

### Competing Interest Statement

The authors have declared no competing interest.

### Summary of Updates

Updated manuscript including the latest upgrade of SAGA software

