## supplementary file for "SAGA (Simplified Association Genome-wide Analyses): a user-friendly Pipeline to Democratize Genome-Wide Association Studies"

**Supplementary Figure 1.** Tools implemented in SAGA and their corresponding workflow.

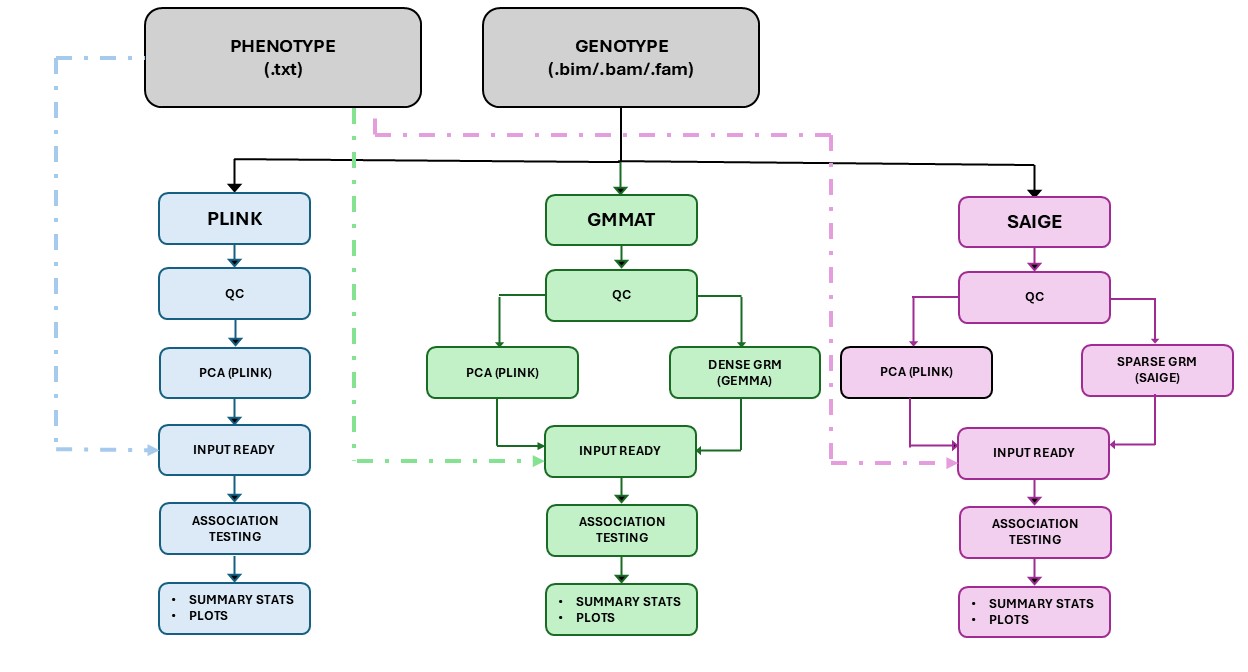

**Supplementary Figure 2.** Manhattan plot, QQ, and SNP density plot for common variant analysis for the ADSP multi-sample using PLINK.

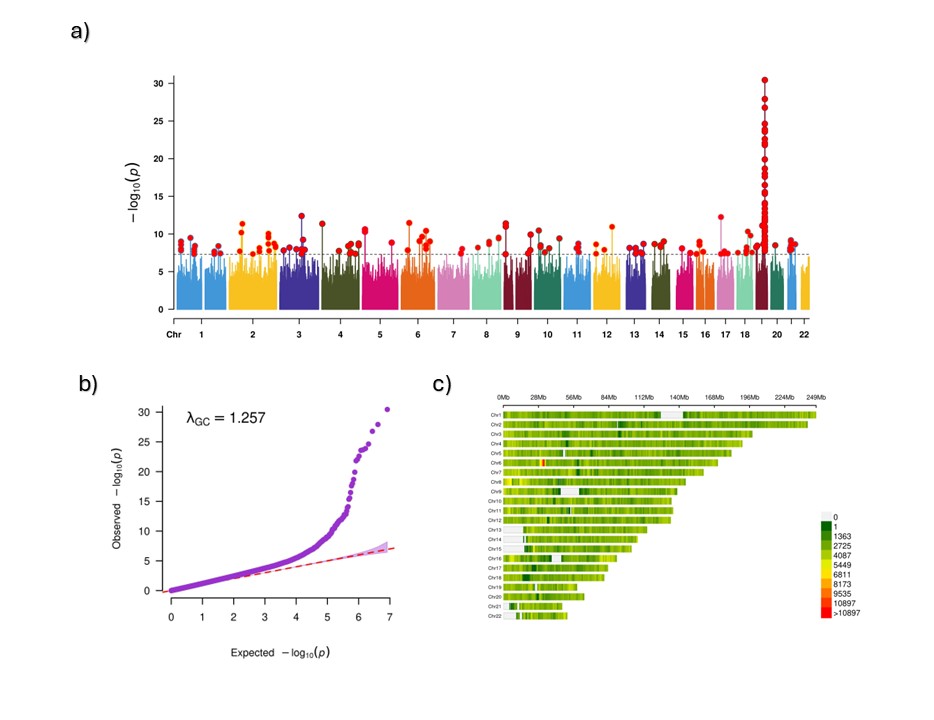

**Supplementary Figure 3.** Manhattan plot, QQ, and SNP density plot for common variant analysis for the ADSP multi-sample using GMMAT.

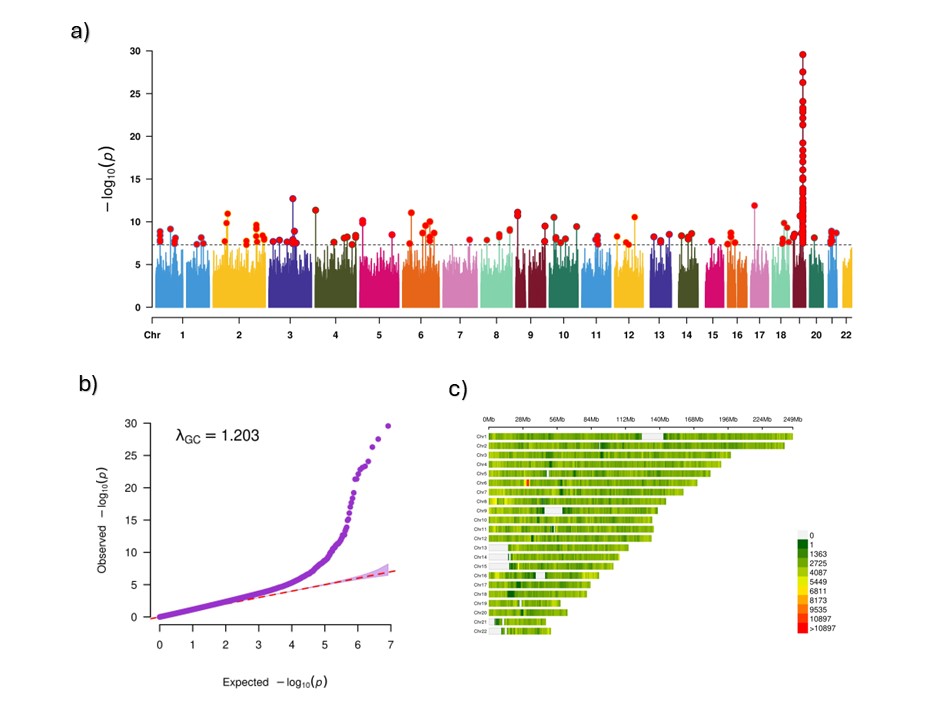

**Supplementary Figure 4.** Manhattan plot, QQ, and SNP density plot for common variant analysis for the PR-1066 cohort using PLINK (a), GMMAT (b), and SAIGE (c).

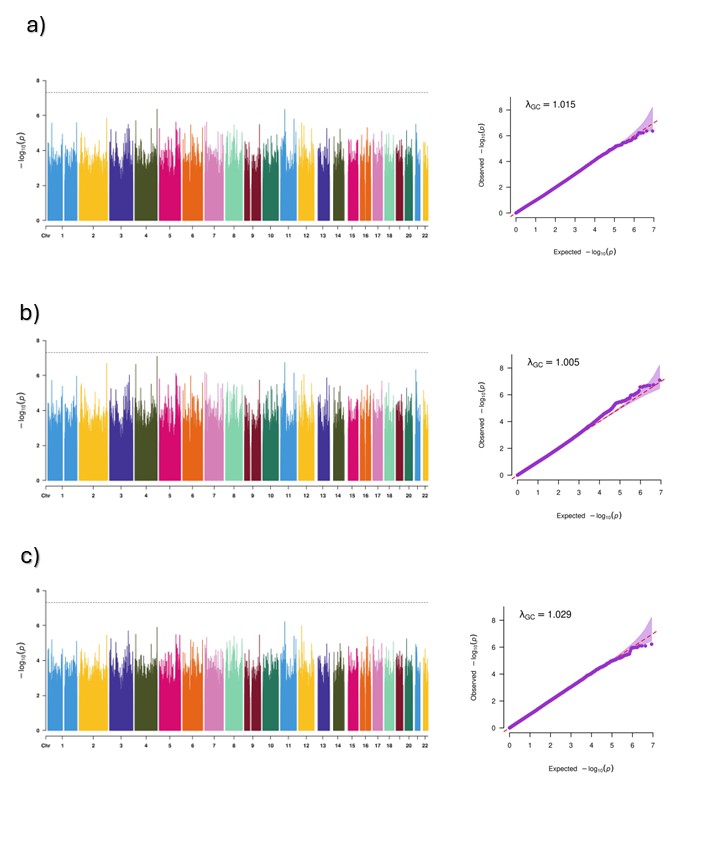

**Supplementary Figure 5.** Manhattan plot, QQ, and SNP density plot for common variant analysis for the ADC cohort using PLINK (a), GMMAT (b), and SAIGE (c).

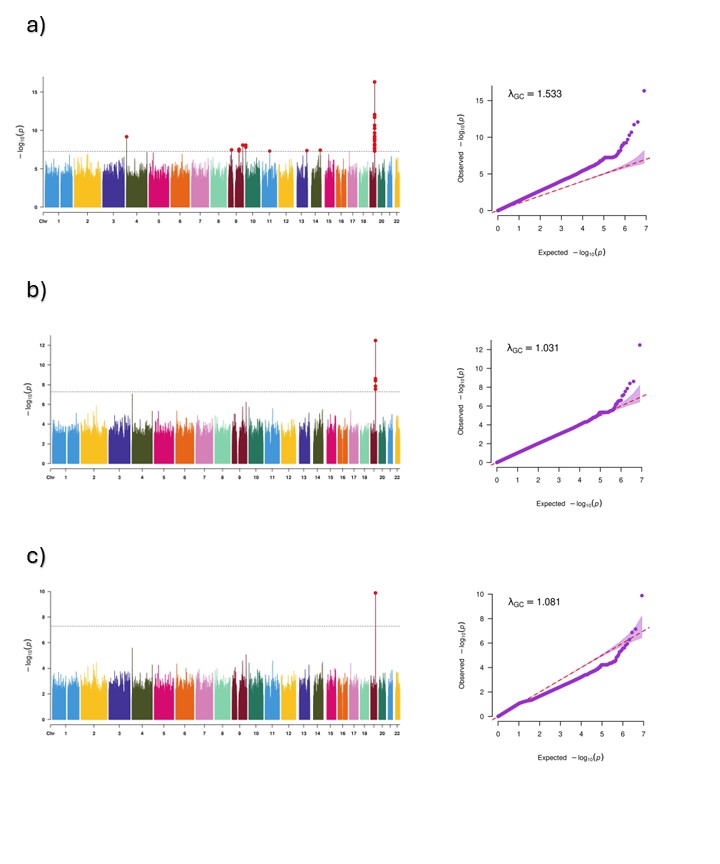

**Supplementary Figure 6.** Manhattan plot, QQ, and SNP density plot for common variant analysis for the PRADI using PLINK (a), GMMAT (b), and SAIGE (c).

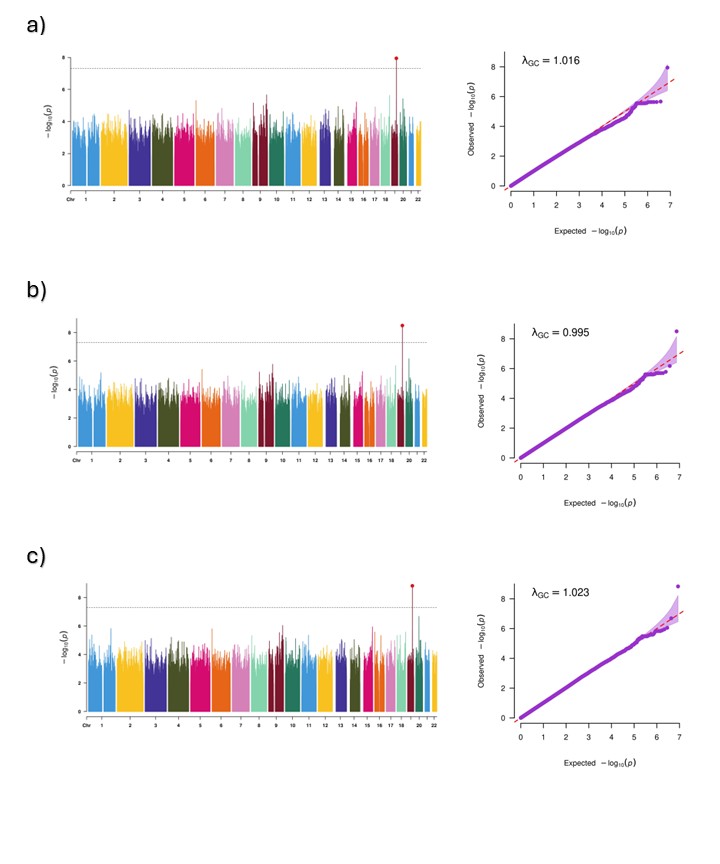

**Supplementary Figure 7.** Manhattan plot, QQ, and SNP density plot for common variant analysis for the TARCC cohort using PLINK (a), GMMAT (b), and SAIGE (c).

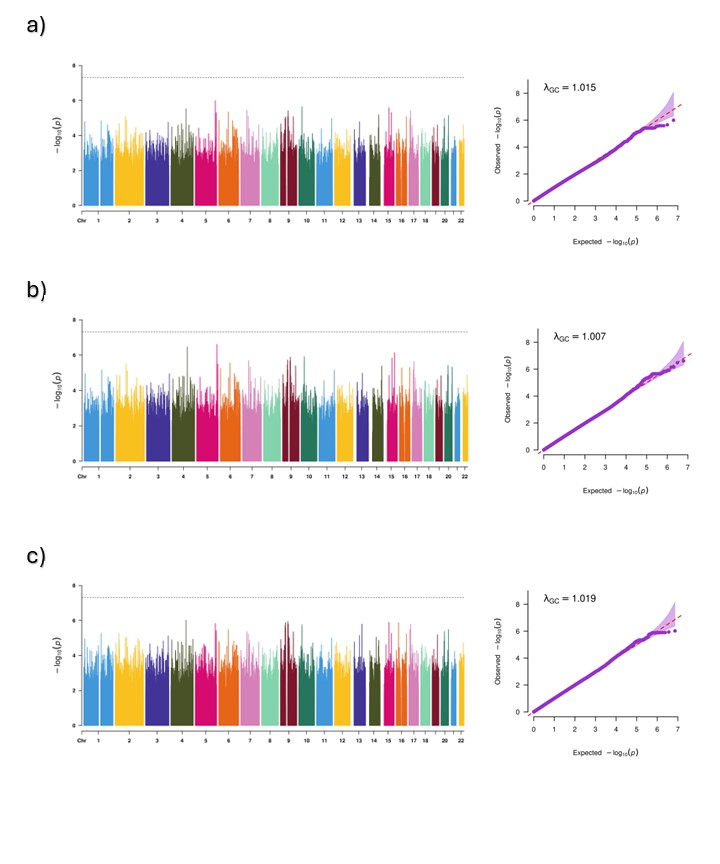

**Supplementary Figure 8.** Manhattan plot, QQ, and SNP density plot for common variant analysis for the NOMAS cohort using PLINK (a), GMMAT (b), and SAIGE (c).

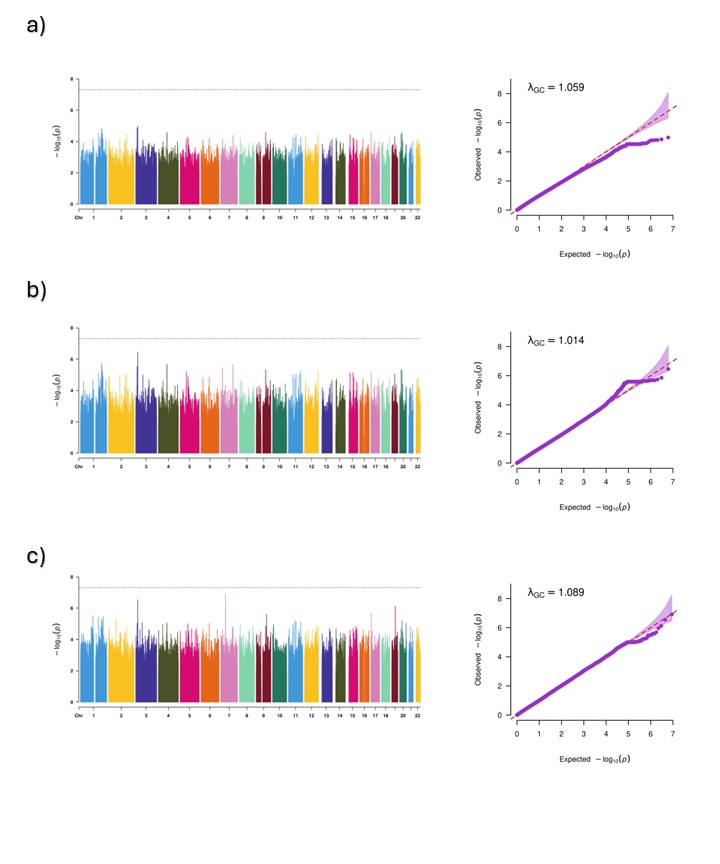

**Supplementary Figure 9.** Manhattan plot, QQ, and SNP density plot for common variant analysis for the PeADI cohort using PLINK (a), GMMAT (b), and SAIGE (c).

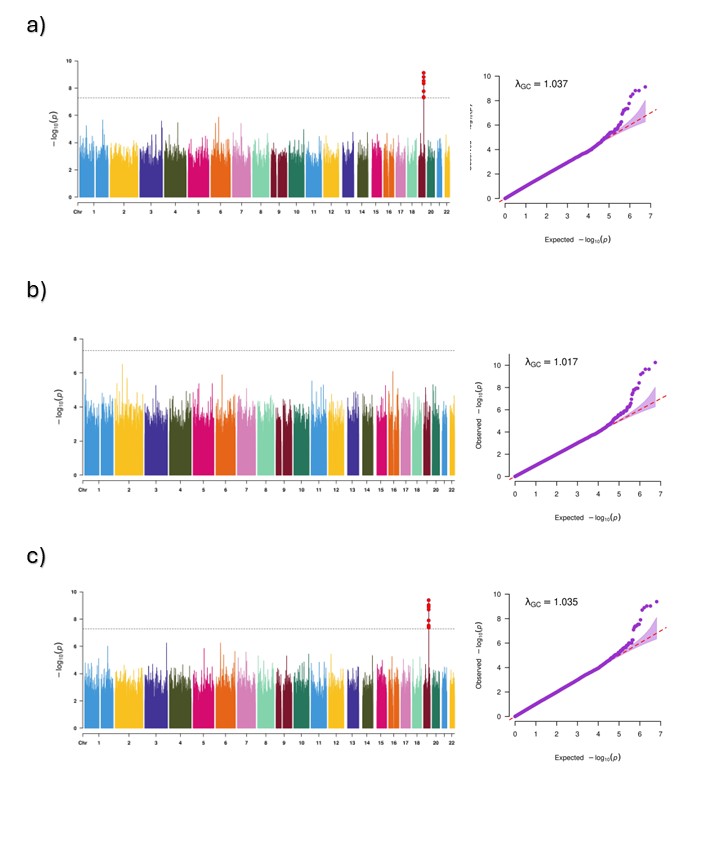

**Supplementary Figure 10.** Manhattan plot, QQ, and SNP density plot for common variant analysis for the WHICAP cohort using PLINK (a), GMMAT (b), and SAIGE (c).

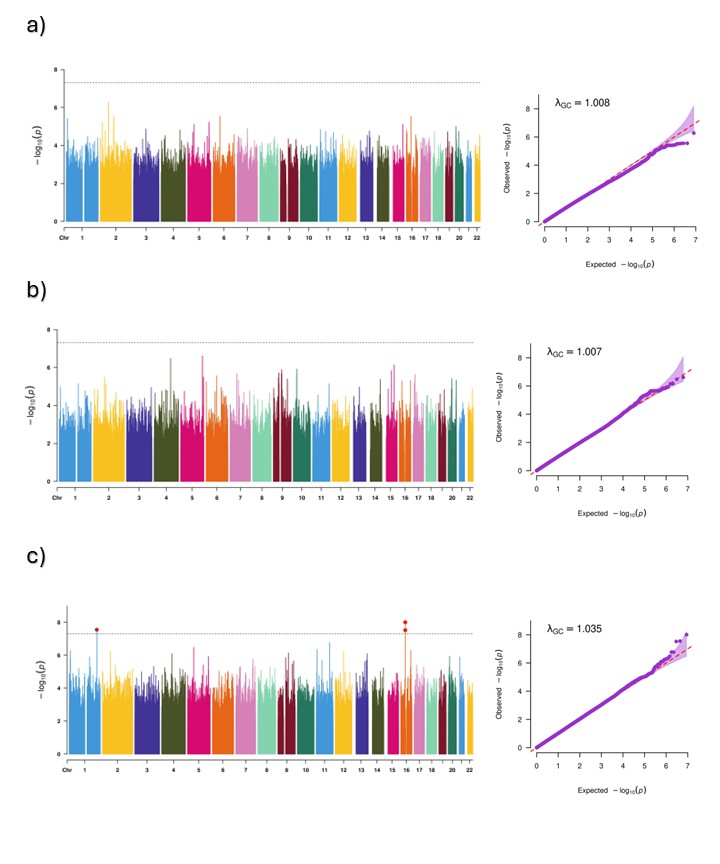

**Supplementary Table 1.** Summary of Quality Control Procedures in SAGA.

| **Purpose** | **QC Step** | **Description** | **Threshold / Criteria** | **Tool Used** | **Output / Action** |
| --- | --- | --- | --- | --- | --- |
| **Association Analysis** | SNP Missingness | Remove variants with excessive missing genotypes | --geno 0.02 (≥98% call rate) | PLINK | Low-quality SNPs removed |
|  | Individual Missingness | Remove samples with high missingness | --mind 0.02 (≥98% call rate) | PLINK | Poor-quality individuals removed |
|  | Minor Allele Frequency (MAF) | Exclude variants with low minor allele frequency | --maf 0.01 (MAF ≥ 1%) | PLINK | Rare variants excluded |
|  | Hardy-Weinberg Equilibrium | Remove variants deviating from equilibrium | --hwe 1e-6 | PLINK | SNPs violating HWE excluded |
|  | **Output Dataset** | Final dataset for GWAS association testing | After stepwise filtering | PLINK | QCed.assoc.{bed,bim,fam} ready for analysis |
| **Kinship Matrix Construction** | SNP Missingness | Remove SNPs with low call rate | --geno 0.02 | PLINK | Low-quality variants removed |
|  | Individual Missingness | Remove samples with high missingness | --mind 0.02 | PLINK | Low-quality individuals removed |
|  | Minor Allele Frequency (MAF) | Keep only common variants to ensure stable GRM | --maf 0.05 (MAF ≥ 5%) | PLINK | Common variants retained |
|  | Hardy-Weinberg Equilibrium | Exclude variants deviating from equilibrium | --hwe 1e-6 | PLINK | SNPs violating HWE removed |
|  | LD Pruning | Remove correlated variants to prevent bias in GRM | --indep-pairwise 50 5 0.2 | PLINK | List of independent SNPs (.prune.in) |
|  | Extract Pruned SNPs | Create dataset of independent, high-quality SNPs | --extract pruned_snps.prune.in | PLINK | QCed.kinship.{bed,bim,fam} for kinship calculation |
|  | **Output Dataset** | Final dataset for GRM/kinship computation | After LD pruning | Clean input for GMMAT/SAIGE kinship module | QCed.kinship.{bed,bim,fam} for kinship calculation |

**Supplementary Table 2.** Code examples.

| **Pipeline** | **Command Example** | **Description / Key Arguments** |
| --- | --- | --- |
| **PLINK-based Association** | bash ./run_pipeline_plink.sh \  full_path_to_geno/geno \  full_path_to_pheno/pheno_binary.txt \  COV1, COV2, COV3, PC1, PC2, PC3 \  PHENO \  binary \  myoutputs | Runs the PLINK association pipeline using binary or quantitative phenotypes.   - geno – genotype files (.bed/.bim/.fam) - pheno_binary.txt – phenotype file - COV1,…,PC3 – comma-separated covariates - PHENO – phenotype column - binary – type of trait - myoutputs – output directory |
| **GMMAT-based Association** | bash ./run_pipeline_gmmat.sh \  full_path_to_geno/geno \  full_path_to_pheno/pheno_binary.txt \  "PHENO ~ COV1 + COV2 + PC1 + PC2 + PC3" \  binary \  myoutputs | Performs mixed-model GWAS using the GMMAT R package.   - geno – genotype files (.bed/.bim/.fam) - pheno_binary.txt – phenotype file - Model syntax follows R formula notation. - binary – type of trait - myoutputs – output directory |
| **SAIGE-based Association** | bash ./run_pipeline_saige.sh \  full_path_to_geno/geno \  full_path_to_pheno/pheno_binary.txt \  COV1,COV2,PC1,PC2,PC3 \  COV1 \  PHENO \ binary \  myoutput | Uses the SAIGE container for large-scale mixed-model GWAS.  Requires Singularity image Saige_1.3.0.sif in SAIGE/.   - geno – genotype files (.bed/.bim/.fam) - pheno_binary.txt – phenotype file - COV1,…,PC3 – List of all covariates; - COV1 – Categorial covariates - PHENO – phenotype column - binary – type of trait - myoutputs – output directory |

**Supplementary Table 3.** Benchmarking of SAGA across sample sizes and SNP densities. Gray color corresponds to results shown in Figure 1 on main text. White non-color cells correspond to results for additional benchmarking using 1000 and 10 000 SNPs.

| **Number of SNPs** | **Sample Size (N)** | **PLINK Runtime (s)** | **GMMAT Runtime (s)** | **SAIGE Runtime (s)** |
| --- | --- | --- | --- | --- |
| 1,000 | 1,000 | 2 | 12 | 19 |
| 1,000 | 5,000 | 20 | 623 | 60 |
| 1,000 | 10,000 | 144 | 4,695 | 276 |
| 10,000 | 1,000 | 3 | 19 | 17 |
| 10,000 | 5,000 | 36 | 1,042 | 60 |
| 10,000 | 10,000 | 201 | 6,466 | 278 |
| **100,000** | **1,000** | **19** | **108** | **103** |
| **100,000** | **5,000** | **181** | **3,218** | **247** |
| **100,000** | **10,000** | **727** | **17,652** | **989** |

1. **Phenotype File**

**Filename:** *pheno_example.txt*

FID	IID	COV1	COV2	PHENO

FAM001	IND001	0	84	1

FAM002	IND002	0	85	0

FAM003	IND003	1	72	1

...

**FID, IID:** Family and individual IDs

**COV1–COV10:** Optional covariates (up to ten)

**PHENO:** Target phenotype (quantitative or binary)

2. **Genotype File**

PLINK files for association testing

- *geno.bed*
- *geno.bim*
- *geno.fam*
